# Assessing aneuploidy with repetitive element sequencing

**DOI:** 10.1101/660258

**Authors:** Christopher Douville, Joshua D. Cohen, Janine Ptak, Maria Popoli, Joy Schaefer, Natalie Silliman, Lisa Dobbyn, Robert E. Schoen, Jeanne Tie, Peter Gibbs, Michael Goggins, Christopher L. Wolfgang, Tian-Li Wang, Ie-Ming Shih, Rachel Karchin, Anne Marie Lennon, Ralph H. Hruban, Cristian Tomasetti, Chetan Bettegowda, Kenneth W. Kinzler, Nickolas Papadopoulos, Bert Vogelstein

## Abstract

We report a sensitive PCR-based assay that can detect aneuploidy in samples containing as little as 3 picograms of DNA. Using a single primer pair, we amplified ∼750,000 amplicons distributed throughout the genome Aneuploidy was detected in 49% of liquid biopsies from a total of 883 non-metastatic cancers of eight different types. Combining aneuploidy with somatic mutation detection and eight standard protein biomarkers yielded a median sensitivity of 80% at 99% specificity.

---

As a result of drastic reductions in costs, whole genome sequencing (WGS) is now commonly used to detect chromosome copy number variations, also known as aneuploidy ^1^. Identifying the presence of aneuploidy has a broad range of diagnostic applications including non-invasive prenatal testing (NIPT) ^2^, preimplantation genetic diagnosis ^3^, evaluation of congenital abnormalities ^4^, and cancer diagnostics ^5^.

Shallow (0.1x-1x) WGS is employed for aneuploidy detection in a large number of commercially available tests ^6^. WGS is typically employed in NIPT, where a relatively high fraction (5-25%) of the total DNA is derived from the fetus ^7^. A companion diagnostic is frequently used to estimate the fetal fraction and NIPT is often not performed when the fraction of fetal DNA is less than 4% ^8, 9^. Sequencing depth becomes a major issue for the assessment of aneuploidy in cell-free DNA from patients with cancer, where the fraction of DNA derived from cancer cells is often much less than 1% of the total input DNA ^10^.

Amplicon-based methods using sequence-specific primers have been proposed as an alternative to WGS for the assessment of aneuploidy ^11-13^. Amplicon-based protocols offer many advantages over WGS (or exome sequencing), including a simpler workflow that does not require library construction, a reduced requirement for input DNA, and a simplified computational analysis. Here, we report a substantially improved amplicon-based approach to detect the presence of aneuploidy, named the Repetitive Element AneupLoidy Sequencing System (REAL-SeqS). Using a single PCR primer pair, REAL-SeqS amplifies ∼750,000 genomic loci with an average size of 88 base pairs spread throughout the genome.

REAL-SeqS introduces six key innovations:

- higher sensitivity with less sequencing than previously reported technologies
- increased spatial coverage throughout the genome, enabling the detection of microdeletions and microamplifications.
- an improved machine-learning-based algorithm for interpreting the data
- reduced requirement for input DNA
- concomitant detection of contaminating cellular DNA in samples of cell-free DNA
- miproved detection of aneuploidy in cell-free DNA.

The FAST-SeqS approach described in Kinde et al was the first aneuploidy detection method to use a single primer pair to amplify numerous repetitive long interspersed nucleotide elements (LINEs) spread throughout the genome ^11^. However, due to the low genomic density of these amplicons (a total of 38,000 across the entire genome), its power to detect focal amplifications and deletions (<5MB) was limited. Additionally, FAST-SeqS amplicons ranged in size from 120-145 bps, which was sub-optimal for assessing cell-free DNA, which has an average size of ∼140 bp. Accordingly, FAST-SeqS was only able to detect aneuploidy in 22% of liquid biopsy samples containing more than 1% of tumor-derived DNA ^14^.

Based on the limitations described above, we attempted to identify a single primer pair that could amplify far more than 38,000 amplicons of a size far less than 120 to 145 bp. To generate a list of candidate primers, we first calculated the frequency of all possible 6-mers (4^6 = 4096) within the RepeatMasker track of hg19. Next, we calculated the frequency of all possible 4-mers (4^4 = 256) within 75 bp upstream or downstream from the 6-mers. Joining the 6-mers with the 4-mers generated 2,097,152 candidate pairs. We narrowed these pairs based on the number of unique genomic loci expected from their PCR-mediated amplification, the average size between the 6-mer and its corresponding 4-mers, and the distribution of these sizes, aiming for a unimodal distribution. This filtering criteria generated 7 potential k-mer pairs, leading to the design of 7 primer pairs that incorporated these k-mer pairs at their 3-ends. Two of these primer pairs (REAL1 and REAL2) outperformed the remaining 5 primers when assessed experimentally by the number of unique loci that were amplified and the size distribution of the amplicons. After further experimental testing of REAL1 and REAL2 on 100 normal samples, the REAL1 primer pair was chosen for the experiments reported herein. The average amplicon size of REAL1 was 88 base pairs (Supplementary Figure 1). Details of the primer selection methods, experimental procedures, new analytic techniques, and work flow diagram are described in Supplementary Text and Supplementary Figure 2.

In the most common form of NIPT, detection of a gain or loss of a chromosome (e.g., chromosome 21 in Down Syndrome) is the goal. We used WGS (Supplementary Table 2), FAST-SeqS (Supplementary Table 3), and REAL-SeqS (Supplementary Table 4) to assess performance on a collection of synthetic samples for DNA admixtures typically encountered in NIPT, i.e., when the fraction of fetal DNA was 5% (pseudocode used to generate synthetic samples Supplementary Figures 3 and 4). To ensure that these comparisons were intrinsic to the sequencing data rather than to the computational algorithm used to analyze the data, we calculated performance using simple z score comparisons (Supplementary Text). We reported results in total reads needed for all three approaches assuming single-end 100 bp reads and accounting for differences in alignment rates and filtering criteria typically used (Supplementary Tables 2, 3, and 4), REAL-SeqS consistently achieved higher sensitivity at lower amounts of sequencing. For example, REAL-SeqS had 98.5% sensitivity (at 99% specificity) for monosomies and trisomies at a 5% cell fraction, while WGS and FAST-SeqS had 93.9% and 81.1% sensitivity respectively (Figure 1A).

**Figure 1.**
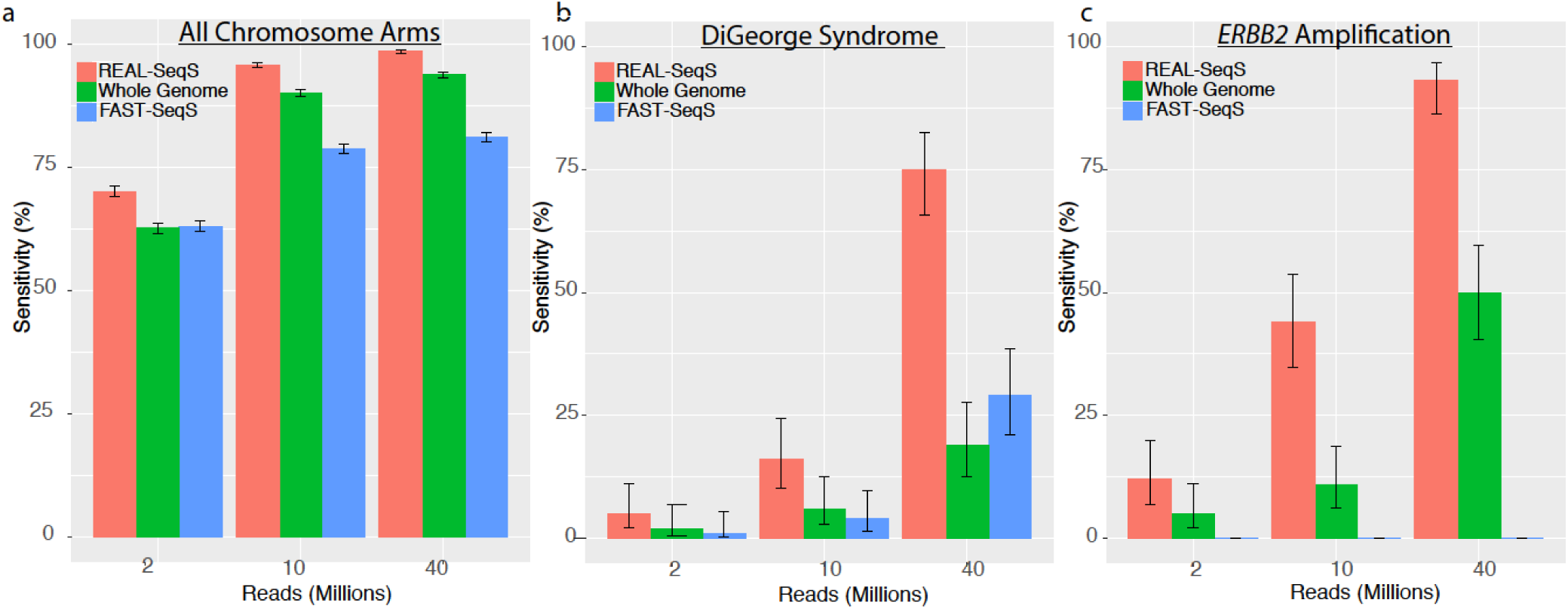
Detection of Aneuploidy using Next Generation Sequencing Technologies. Sensitivities were calculated using a threshold at 99% specificity. Error bars represent 95% confidence intervals. (a) Comparison of sensitivity for monosomies and trisomies across all 39 non-acrocentric chromosome arms at 5% cell fraction. (b) Comparison of sensitivity for the 1.5 Mb DiGeorge deletion on 22q at 5% cell fraction. (c) Comparison of sensitivity for a 20 copy *ERBB2* focal amplification at 1% cell fraction.

Another important aspect of assays for copy number variation is the detection of relatively small regions which are deleted or amplified. For example, the DiGeorge Syndrome deletions are often as small as 1.5 Mb ^15^. For a 5% deletion-containing cell fraction, REAL-SeqS had 75.0% sensitivity for the 1.5 Mb DiGeorge deletion (at 99% Specificity) while WGS and FAST-SeqS had 19.0% and 29.0% sensitivity, respectively (Figure 1B).

The detection of amplifications, such as those on *ERBB2* in breast cancer, are critical for deciding whether patients should be treated with trastuzumab or other targeted therapies. Following the same protocol as above, we generated synthetic samples with focal amplifications of the ∼42 Kb ERBB2 gene (20 copies) for WGS, FAST-SeqS, and REAL-SeqS. REAL-SeqS could detect such amplifications in the synthetic samples with significantly less sequencing than could WGS or Fast-SeqS. For a 1% cell fraction, REAL-SeqS had a 91.0% sensitivity while WGS had 50.0% (Figure 1C). FAST-SeqS did not have enough spatial coverage in this genomic region to detect *ERBB2* amplifications.

Reliably detecting aneuploidy in only a few pg of DNA is necessary for preimplantation diagnostics as well as forensic applications. In preimplantation diagnosis, a few cells picked from a blastocyst are used to assess copy number variations, such as those responsible for Down Syndrome. To test the limit of detection of REAL-SeqS with respect to input DNA, we analyzed trisomy 21 samples at input DNA concentrations ranging from 3-34 pg (Supplementary Figure 5 and Supplementary Data 2). Trisomy 21 was detected in all these samples, even those from 3 pg of DNA, representing half of a diploid cell. No chromosome arms other than chromosome 21 were found to be aneuploid in the Trisomy 21 samples. No chromosome arms, including chromosome 21, were found to be aneuploid in the euploid controls used in these experiments. The reduced requirement for input DNA also enables retrospective testing of samples from biobanks for either aneuploidy or identification purposes (using SNPs within the amplified repeated sequences; see Supplementary Material).

Plasma “cell-free” DNA is often contaminated with DNA that has leaked out of leukocytes, either during phlebotomy or preparation of plasma. This contaminating leukocyte DNA can reduce the sensitivity of aneuploid testing for plasma samples because leukocytes are not derived from either fetal cells (in NIPT) or cancer cells (in liquid biopsies). Leukocyte DNA (gDNA) has an average size of >1000 bp while cell-free plasma DNA has an average size of < 160 bp. DNA size impacts PCR efficiency and long amplicons may not be present in cfDNA. REAL-SeqS enables the detection of leukocyte DNA contamination by virtue of the differently-sized amplicons generated with REAL1 primers. We identified 1241 amplicons typically present in gDNA but not cfDNA (Supplementary Data 3). Reads at these amplicons thereby indicates leukocyte contamination in plasma samples (Supplementary Data 3). Through mixing of leukocyte DNA with cell-free plasma DNA, we were able to demonstrate that samples containing >4% of leukocyte DNA could be detected with REAL-SeqS (Supplementary Table 5).

DNA from cancer cells is shed into the bloodstream, fostering the analysis of cell-free DNA in plasma (“liquid biopsies”) to detect the presence of cancers. Several features of cancer DNA, including point mutations, aberrant DNA methylation, and aneuploidy have been used to assess liquid biopsies. Because aneuploidy is a feature of virtually every cancer type (>90%), it is well-suited for this purpose ^14, 16^.

REAL-SeqS was used to detect aneuploidy in cell-free plasma DNA from 883 patients harboring surgically resectable cancers of 8 different cancer types (ovary, colorectum, esophagus, liver, lung, pancreas, stomach, and breast). Each plasma sample was given a REAL-SeqS score based on a machine-learning-based algorithm described in the Supplementary Text. Aneuploidy was scored in the samples from cancer patients at a threshold of 99% specificity derived from the analysis of 1348 plasma samples from healthy individuals (Supplementary Data 4 and 5). The plasma samples from cancer patients had previously been analyzed for somatic point mutations and small insertions or deletions using a sensitive mutation detection technique based on 61 genomic regions that are frequently altered in cancer ^10^. Mutations in the plasma samples were also scored at a threshold of 99% specificity.

Overall, we found that aneuploidy was detected more commonly than mutations (49% and 34% of 883 samples, respectively) in plasma samples from cancer patients (P < 10^−20^, one sided binomial test) With respect to tissue type, aneuploidy was detected more commonly than mutations in samples from patients with cancers of the esophagus, colorectum, pancreas, lung, stomach, and breast, (all P-values <0.01), less commonly in ovary (P=0.048), and equally commonly in liver cancer (Figure. 2A). Importantly, aneuploidy was detected in 242 (42%) of plasma samples in which mutations were not detected, and conversely, mutations were detected in 112 (25%) of samples in which aneuploidy was not detected. Higher amounts of tumor DNA were associated with higher sensitivities. Aneupoidy was detected in 89 of 94 (95%) samples of high ctDNA content (somatic mutation allele frequency >1%). Mutations in the plasma originating from clonal hematopoiesis of indeterminant potential (CHIP), rather than from cancer cells, has confounded previous analyses of mutations in cell-free DNA. This confounder was mitigated with aneuploidy detection; 0 of 17 samples that had CHIP mutations were positive for aneuploidy. We also tested leukocyte DNA from 18 patients whose plasma samples scored positive for aneuploidy with REAL-SeqS; only one of these leukocyte samples was aneuploid as assessed by REAL-SeqS.

**Figure 2.**
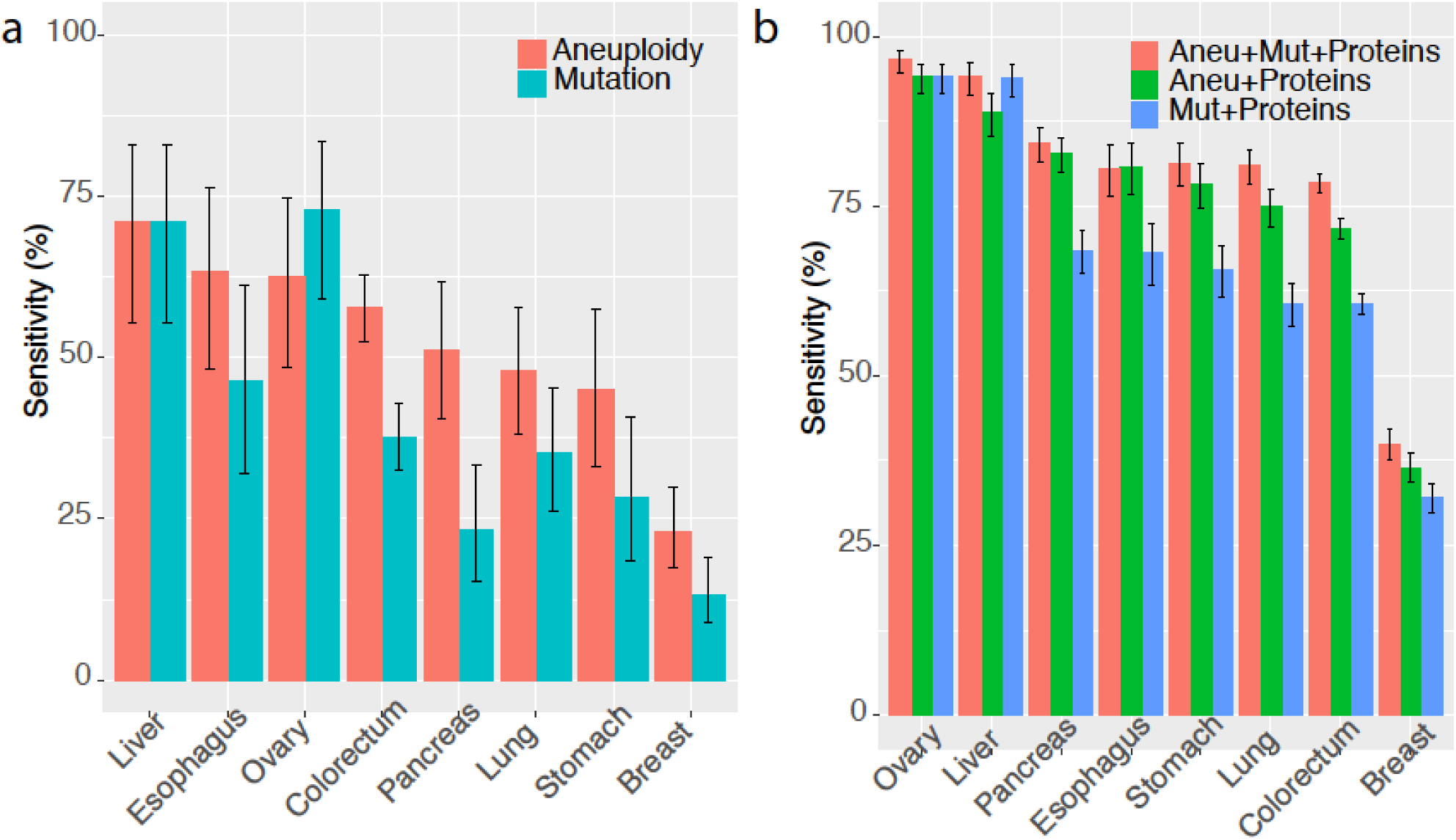
Detection of cancer in liquid biopsies from samples with non-metastatic cancers of eight different types. Sensitivities were calculated using a threshold at 99% specificity. Error bars represent 95% confidence intervals. (a) Comparison of aneuploidy status as calculated by REAL-SeqS to somatic mutations status. (b) Comparison of different multi-analyte tests. Three different multi-analyte tests evaluated sensitivity with and without the inclusion of aneuploidy status with somatic mutation status and 8 standard protein biomarkers.

Either mutations or aneuploidy were detected in 551 of the 883 plasma samples (62%). This performance is likely an underestimate of what could be achieved: more mutations might have been detected if more amplicons were sequenced and additional aneuploidy might have been identified at a greater sequencing depth. However, in practice there must be a balance between sensitivity and cost, thus limiting the amount of sequencing that can be performed in a screening setting.

Finally, we evaluated the ability of a multi-analyte test, combining protein biomarkers for cancer with aneuploidy and mutations in the cohort described above (detailed methods in Supplementary Text). Eight protein biomarkers were evaluated in these 883 cancer samples as previously described ^10^. We scored plasma samples using a logistic regression model maintaining an aggregate specificity of 99% (Supplementary Data 5, 6, and 7). In the plasma samples of patients harboring cancers of seven tissue types (liver, ovary, pancreas, esophagus, stomach, colorectal, and lung), the median sensitivity was 80% (range 77% to 97%), while in breast cancers, it was 38% (Figure 2B).

In summary, REAL-SeqS is exceedingly simple to perform, requires only a single primer pair, and is relatively sensitive and cost effective (∼$110 per assay). We anticipate that it will be used to assess aneuploidy in a variety of clinical contexts.

## Acknowledgements

This work was supported by the Lustgarten Foundation for Pancreatic Cancer Research, The Virginia and D.K. Ludwig Fund for Cancer Research, The Commonwealth Fund, Burroughs Wellcome Career Award for Medical Scientists, The Honorable Tina Brozman Foundation, Gray Foundation, Susan Wojcicki and Dennis Troper, the Rolf Foundation, The Templeton Foundation, and NIH grants T32-GM007309, NCI grants U01CA230691-01, P50CA228991, U01CA200469, R37 CA230400-01, CA62924, CA210170.

## Author Contributions

C.D., N.P., K.W.K., and B.V. designed research; C.D., N.P., K.W.K., and B.V. performed research; C.D., J.D.C., J.P., M.P., J.S., N.S., L.D., R.E.S., J.T., P.G., M.G., C.L.W., T.L.W., I.M.S., R.K., A.M.L., R.H.H., C.B.,C.T., N.P., K.W.K., and B.V. contributed new reagents/analytic tools; C.D. and B.V. analyzed data; and C.D. and B.V. wrote the paper.

