## Supplementary Text for "Assessing aneuploidy with repetitive element sequencing"

**Supplementary Figures**

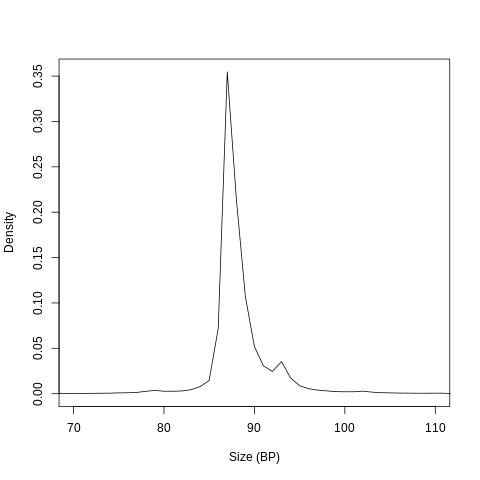

**Supplementary Figure 1**. Distribution of amplicon sizes for REAL-SeqS

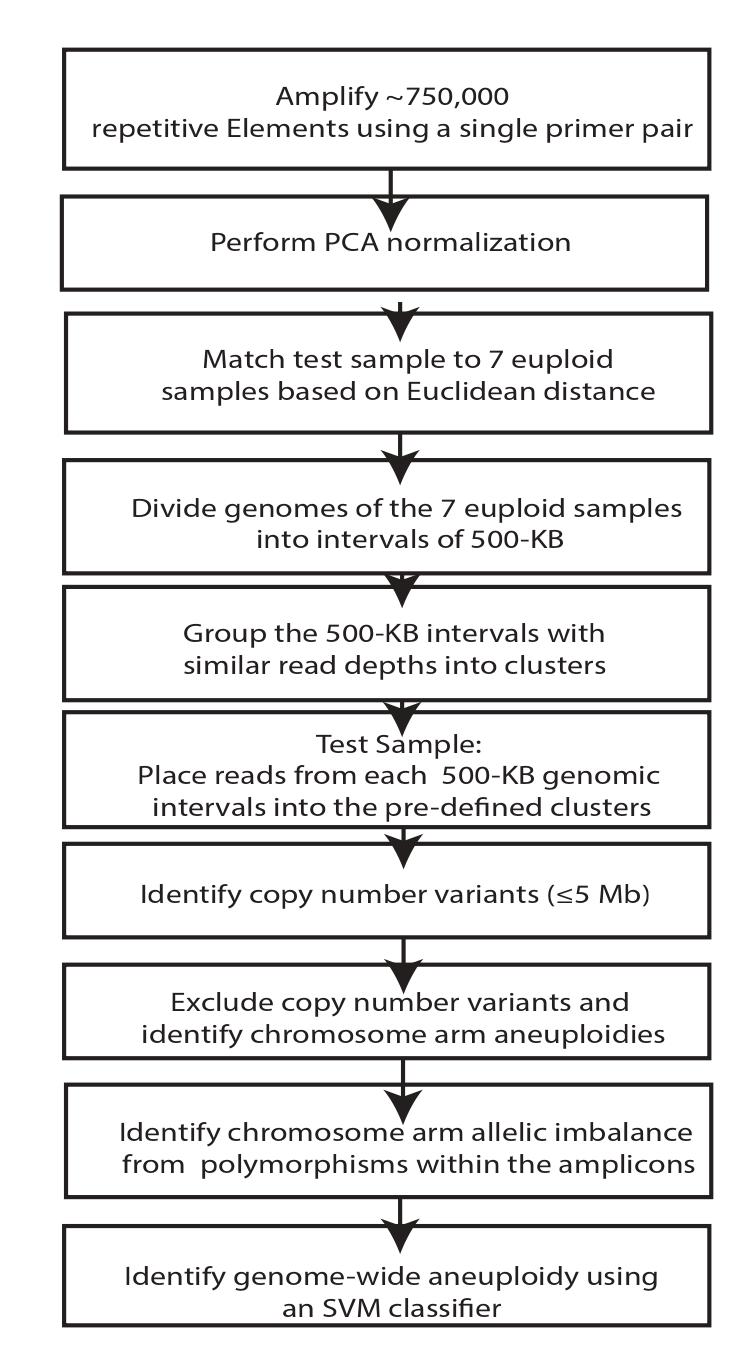

**Supplementary Figure 2.** Overview of the work the REAL-SeqS workflow.

Synthetics with one arm alterations

N <- number of normal samples used

f <- desired neoplastic cell fraction

r <- chromosome arm (1p..22q)

r_type <- alteration type (gain or loss)

rho <- desired unique read depth of synthetic sample

j<-desired number of repeats

For j=1:5

For i=1:N

s <- sample[i]

s_fraction<-vector that contains the fraction of reads that map to each of the 39 chr arm

for sample s

For f in 1:F

For r in 1p:22q

For r_type in gain loss

h <- new synthetic sample

t <- copy of s

t_fraction<- vector that contains the fraction of reads that map

to each of the 39 chr arm for sample t

if r_type== gain

t_fraction[r]<-t_fraction[r]*3/2 ###trisomy on arm r

else (r_type == loss)

t_fraction[r]<-t_fraction[r]*1/2###monosomy on arm r

Endif

h_normal<-weighted random select (1-f)*rho where the

weights are r_fraction

h_aneuploid<-weighted random select f*rho where the weights

are t_fraction

h<-h_normal+h_aneuploid

End

End

End

End

End

**Supplementary Figure 3.** Pseudocode to generate the synthetic trisomy and monosomy samples used for the comparison of whole genome sequencing, FAST-SeqS, and REAL-SeqS.

N <- number of normal plasma samples used

d <- desired degree of aneuploidy (number arms altered)

f <- desired neoplastic cell fraction

r <- chromosome arm (1p..22q)

r_type<-arm alteration type (gain or loss)

p(r) <- probability that an arm is gained or lost in cancer (Estimated from ^1^)

rho <- desired unique read depth of synthetic sample (10M default)

j<- desired number of repeats

For j=1:5

For i=1:N

s <- sample[i]

U <- get reads from s

For f in 1:F

h <- new synthetic sample

For d in (10,15,20) #desired numbers of altered arms

t <- copy of s

For a in 1:d #select alteration types and spike-in to t

r, r_type <- weighted random selection where the weights [p(r)] are the likelihood that an arm is gained or loss in cancer

u_all <- get all Reads that that map to chr arm r

if r_type== gain

t <- add 50% of u_all reads

else (r_type == loss)

t <- subtract 50% of u_all reads

Endif

h_normal<-randomselect (1-f)*rho reads from s

h_aneuploid<-randomselect f*rho reads from t

h<-h_normal+h_aneuploid

End

End

End

End

End

**Supplementary Figure 4**. Pseudocode to Generate Synthetics with multiple arm alterations that were used in the Genome Wide Aneuploidy SVM training set.

**Supplementary Text**

**Sample Information**

Plasma was purified from 1348 healthy individuals and 883 patients with cancer using Qiagen kitcatalog #927255. All individuals participating in the study provided written informed consent after approval by the institutional review board at the patients' participating institutions.

**Primer Development**

As noted in the main text, we first calculated the frequency all possible 6-mers (4^6 = 4096) within the RepeatMasker track for hg19. Next, we calculated the frequency of all possible 4-mers (4^4 = 256) within 75 bp upstream or downstream from the 6-mers. Joining the 6-mers with the 4-mers generated 2,097,152 candidate pairs. We narrowed these pairs based on the number of unique genomic loci expected from their PCR-mediated amplification, the average size between the 6-mer and its corresponding 4-mers, and the distribution of these sizes, aiming for a unimodal distribution. This filtering criteria generated 7 potential k-mer pairs that were each used to design primer pairs for PCR. Nucleotides were added to each kmer pair to generate functional primer pairs (Supplementary Table 1) with a 4-mer at the 3'-end of one primer and a 6-mer at the 3'-end of the other primer. Nucleotides upstream of the 4-mer and 6-mers within the primers were then added, informed by the most common nucleotides at the predicted amplicons and their predicted melting temperatures; optimally, all amplicons generated with a specific primer pair should have identical melting temperatures. Two of the 7 primer pairs (REAL1 and REAL2) outperformed the remaining 5 primers in pilot tests, as assessed by the number of unique loci that could be amplified and the size distribution of the amplicons. After further experimental testing on 100 samples, REAL-1 was chosen for all of the other experiments described in the text.

Bowtie2 was used to align reads of the amplicons generated with each of the 7 primer pairs to the human reference genome assembly GRC37 ^2^. With REAL-1 primers, an average of 51.1% of the total reads could be uniquely aligned and the average amplicon size is 88bp (Supplementary Figure 1). REAL1 was theoretically able to amplify up to 745, 184 repetitive elements (Supplementary Data 1) but the average sample contained an average of 350,000 repetitive elements. We believed there were several potential reasons for the discrepancy between the potential number and the actual observed number of amplicons in plasma samples: 1)Polymorphisms within the sequences may cause misalignment and result in “missing amplicons;” 2) Polymorphisms within the primers may not amplify. 3) Each amplicon has a different PCR efficiency. Low efficiency amplicons may be outcompeted during PCR and not be present. 4) Smaller DNA fragments are preferentially amplified and long amplicons (>100 bp) may not be amplified. 5) Long amplicons maybe absent in cell free DNA due to the small sizes of the DNA fragments in cell free DNA. 6) The amount of sequencing used for these samples may not be high enough to observe every amplicon especially those with low PCR efficiencies. 7) Finally, some repetitive elements may not be present in every individual. Within REAL1’s amplicons, we identified 52,762 polymorphisms. The average number of heterozygous sites in our cohort our 1348 normal plasmas and 883 plasmas from cancer patients individuals was 2,200 and these sites could be used to measure allelic imbalance, genetically identify samples, and determine whether samples had been accidentally mixed together. Scripts for allelic imbalance and sample identity are in a github repository and available upon request. Using the same SNPs, synthetic experiments were used to estimate that sample mixing could be detected when the amount of sample 1 DNA was >4% of the amount of sample 2 DNA in a given mixture.

**Experimental Protocol**

PCR was performed in 25 uL reactions containing 7.25 uL of water, 0.125 uL of each primer, 12.5 uL of NEBNext Ultra II Q5 Master Mix (New England Biolabs cat # M0544S), and 5 uL of DNA. The cycling conditions were: one cycle of 98°C for 120 s, then 15 cycles of 98°C for 10 s, 57°C for 120 s, and 72°C for 120 s. For experiments with plasma, the amount of DNA in 5 uL was 0.14 ng. A second round of PCR was then performed to add dual indexes (barcodes) to each PCR prior to sequencing. The forward and reverse primers used for the second round of PCR are listed in Supplementary Table 1. The second round of second round of PCR was performed in 25 uL reactions containing 7.25 uL of water, 0.125 uL of each primer, 12.5 uL of NEBNext Ultra II Q5 Master Mix (New England Biolabs cat # M0544S), and 5 uL of DNA containing 5% of the PCR product from the first round. The cycling conditions were: one cycle of 98°C for 120 s, then 15 cycles of 98°C for 10 s, 65°C for 15 s, and 72°C for 120 s. Amplification products from the second round were purified with AMPure XP beads (Beckman cat # XXX), as per the manufacturer's instructions, prior to sequencing. Each sample was amplified in eight independent PCRs in the first round. Each independent PCR was then re-amplified using index primers in the second PCR round. The sequencing reads from the 8 replicates were summed in the bioinformatic analysis..

All oligonucleotides were purchased from IDT (Coralville, Iowa). Massively parallel sequencing was performed on an Illumina HiSeq 4000. During the second round of PCR, degenerate bases at the 5’ end of one of primers were used as molecular barcodes to uniquely label each DNA template molecule ^3^. This ensured that each DNA template molecule was counted only once. In all instances in this paper, the term “reads” refers to uniquely identified reads. Depending on the experiment, each read was sequenced between 1 and 3 times. An average of 13.2 million reads per sample (IQR 7.9M to 15.2M) were assessed.

**Real-SeqS Bioinformatic Analysis**

The Within-Sample Aneuploidy DetectiOn (WALDO) approach was developed to detect the presence of aneuploidy in amplicon sequencing reads ^1^. Unlike most WGS approaches that assess aneuploidy, WALDO does not compare normalized read counts in a test sample to the normalized read counts in a panel of normal samples. Direct comparisons are subject to batch effects and can introduce new artifacts ^4^. WALDO attempts to mitigate problematic batch effects using a within sample comparison. We tailored this approach for our REAL-SeqS assay and made several analytical improvements (Supplementary Figure 2). The major modifications included a new normalization step, a new way to call small copy number changes of indeterminate length, and an improved way to detect genome-wide aneuploidy, as described below. These analytical improvements coupled with the increased genomic density of amplicons achieved with REAL1 primers enabled greater sensitivity as well as the detection of focal amplifications and deletions less than 1 Mb in size, which was not possible with FAST-SeqS.

**Normalization**

We employed a new method of normalization that reduced the amount of variability between samples. In this normalization, a principal component analysis (PCA) was first performed on sequencing data from the controls. PCA reduced the number of 500 kb genomic intervals from n=5,344 to a more manageable number of dimensions. Using the PCA coordinates of the controls, we modeled whether a particular 500kb interval will be amplified more or less efficiently in future samples based on their PCA coordinates.

$$Correction Factor for 500kb Interval_{i}=\beta_{oi}+\beta_{1i}*PCA_{1}+\beta_{2i}*PCA_{2}+\beta_{3i}*PCA_{3}+\beta_{4i}*PCA_{4}+\beta_{5i}*PCA_{5}$$

For each test sample, we projected the sample into PCA space and calculate the correction factor for each 500kb interval as function of its PCA coordinates. After applying the correction factor to each 500 kb genomic interval, the test sample was matched to 7 control samples based on the closest Euclidean distance of the 500 kb intervals. Scripts for allelic imbalance and sample identity are in a github repository and available upon request.

**Copy Number Analysis of Indeterminate Length**

The original WALDO method required the specification of a particular genomic region of interest (usually an entire chromosome arm) and then calculating the statistical significance of the desired region. For REAL-SeqS, we incorporated the ability to detect copy number variants of indeterminate length. To do so, we calculated the log ratio of the observed test sample and WALDO predicted values from every 500 kb interval across each chromosomal arm. Using the log ratio, we applied a circular binary segmentation algorithm ^5^ to find copy number variants throughout each chromosome arm. Any copy number variant ≤ 5Mb in size was flagged. Before calculating the statistical significance across each chromosome arm, these flagged CNVs were removed. Because we were interested in chromosomal abnormalities affecting a large part of a chromosome arm, these small CNVs were not of general interest to us. However, the small CNVs could be used to assess microdeletions or microamplifications, such as those occurring in DiGeorge Syndrome (chromosome 22q11.2 or in breast cancers (chromosome 17q12).

**Genome Wide Aneuploidy**

A two-class support vector machine (SVM) ^6^ was trained to discriminate between euploid samples and aneuploid samples. The training set contained a negative class of 1348 presumably euploid plasma samples from normal individuals containing at least 2.5 M reads and 635 aneuploid samples. The aneuploid class contained a mixture of synthetic and actual aneuploid samples. SVM training was done with the e1071 package in R, using radial basis kernel and default parameters ^7^. Each sample had 39 Z-score features, representing chromosome arm gains and losses. During training, the positive class was randomly sampled so that the positive class was 10% the size of the negative class. The positive class was randomly sampled at a ratio of two real samples to one synthetic sample. Ten iterations of this procedure were performed. The final genome wide aneuploidy score was the average of the raw svm score across the 10 iterations.

**Sample Exclusion Criteria**

To ensure that all samples included in the results section of paper were of high quality, we developed several exclusion criteria. 1) Samples with less than 2.5M reads were excluded. 2) Samples with sufficient evidence of contamination were excluded. To be labeled as contaminated, the sample had to have at least 10 significant allelic imbalanced chromosome arms (z score >= 2.5) and fewer than ten significant chromosome arms gains or losses (z >= 2.5 or z<= -2.5). Allelic imbalance is determined from SNPs, while gains or losses were assessed through WALDO ^1^. As determined through mixing experiments, a relatively large number of allelic imbalanced chromosome arms in the absence of a large number of gains or losses indicated contamination of the sample with DNA from another individual. 3) In plasma analyses, we excluded samples in which more than 8.5% of the amplicons were larger than 94 bps (50 base pairs between the forward and reverse primers) bp. Such samples were likely to be contaminated with leukocyte DNA 4) Samples outside the dynamic range of our assay, as defined by the equation below.

$$QC Dynamic Range Metric=\sum_{i}^{2q,3q,4q,5q,6q,8q,13q} \frac{Reads on chr_{i}}{\sum_{j=1}^{39} Reads on chr_{j}}$$

The distribution of this metric has long tails. We selected >0.2450 and 0.2320 as dynamic range that we could evaluate cutoffs. 5) Plasma samples with known aneuploidy in the leukocytes of the same patients; such patients were assumed to have Clonal Hematopoiesis of Indeterminate Potential (CHIP) or congenital disorders.

**Generation of synthetic aneuploidy samples.**

We first selected data from 84 presumably euploid plasma samples, each containing at least 10 million reads, and each derived from the DNA of normal WBCs. Synthetic aneuploid samples were created by adding (or subtracting) reads from several chromosome arms to the reads from these normal DNA samples. We added or subtracted the reads from 1, 10, 15, or 20 chromosome arms to each sample. The additions and subtractions were designed to represent neoplastic cell fractions ranging from 0.5% to 1.5% and resulted in synthetic samples containing exactly ten million reads. The reads from each chromosome arm were added or subtracted uniformly. For example, when we modeled five chromosome arms that were lost, each was lost to the identical degree and we did not incorporate tumor heterogeneity into the model. Furthermore, we did not create synthetic samples containing more than three of any chromosome arm; e.g. 4 copies of chromosome 3p. This simplified approach did not comprehensively cover all biologically plausible aneuploidy events. However, limiting the possible combinations of altered arms made sample generation computationally tractable, and the resulting support vector machine worked well in practice. The synthetically generated samples in which reads from only a single chromosome arm were added or subtracted enabled us to estimate the performance of WALDO when only a single chromosome arm of interest was gained or lost. The pseudocode to generate synthetic samples is in Supplementary Figure 3 and Supplementary Figure 4.

**Comparison to Various Massively Parallel Sequencing Technologies**

Ten normal plasma samples were identified for whole genome sequencing, FAST-SeqS, and REAL-SeqS.

We selected 10 publicly available normal plasma samples that had whole genome sequencing ^8^. Each of the 10 samples had the equivalent of 1 HiSeq lane of sequencing (~144,000,000) reads. The authors performed bioinformatic filters to remove highly polymorphic locations ^9^ and common insertions/deletions ^10^. Reads on the sex chromosomes, contigs, and acrocentric chromosome arms were dropped. The fraction of reads for each of the 39 the non-acrocentric chromosome arms were calculated from the remaining reads. The fraction of reads for each of the 39 non-acrocentric autosomal chromosome arms was reported in their publication and reproduced in Supplementary Table 2.

We then selected 10 normal samples with FAST-SeqS with an average of 10M reads and 10 normal samples with REAL-SeqS with an average of 28M reads. The fraction of reads on each chromosome arm was reported in Supplementary Table 3 and Supplementary Table 4. Synthetic aneuploidy samples were generated at 5% cell fraction to represent the amount of fetal DNA often present in noninvasive prenatal testing (NIPT) in order to compare whole genome sequencing, FAST-SeqS, and REAL-SeqS. We evaluated trisomies and monosomies for each of the 39 chromosome arms as well as the 1.5MB DiGeorge deletion (22:19009792-20509792). Next, we generated synthetic focal amplifications of ERBB2 (17:37819167-37911679 20 copies) at 1% cell fraction to represent the amount of tumor DNA present in later stage liquid biopsies. FAST-SeqS does not have the spatial genomic density to cover the region of interest of ERBB2 and could not be evaluated for those synthetic samples.

Aneuploidies were generated at various sequencing depths of 2M, 10M, 40M reads. The statistical significance was calculated using a simple z score. Sensitivities were calculated based on Z>2.575 and Z<-2.575 (alpha=0.01).

$$z_{chr@Depth}=\frac{Observed_{@Depth}-\mu_{normal panel}}{\sigma_{normal panel @Depth}}$$

**Reduced Requirements for DNA Input**

A major advantage of amplicon sequencing is the reduced requirement for input DNA. To test whether REAL-SeqS could reliably detect aneuploidy with less than one genome equivalent, we evaluated trisomy 21 samples and normal euploid DNA at various low input DNA amounts (3-225pg). The concentration of input DNA for these samples was estimated based on the number of reads for each sample. The relationship of reads to DNA was based on negative controls (water wells with no DNA) and the known concentration of the euploid control (y=7e-5*x-0.5196). Trisomy 21 was detected in samples (z>5). No other arms were aneuploid in the T21 samples. No arms were aneuploid in the controls (Supplementary Data 2).

**Detection of leukocyte DNA in cell free DNA**

Leukocyte DNA (gDNA) has an average size of >1000 bp while cell-free plasma DNA has an average size of < 160 bp. DNA size impacts PCR efficiency and long amplicons may not be present in cfDNA. REAL-SeqS enables the detection of leukocyte DNA contamination by virtue of the amplicons generated with REAL-1 primers.

We selected 50 plasma samples aand 50 gDNA samples. We found 1241 amplicons (Supplementary Data 3) that were not present in the plasma samples (< 5 reads across all samples) but were present in the gDNA samples (>1000 reads across all samples).

We mixed DNA from leuokocytes into cell free DNA from a euploid plasma sample at various dilutions ranging from 4% to 54%. The fraction of reads mapping to the 1241 amplicons in the contaminated samples were more than 3 times higher than the euploid sample (Supplementary Table 5).

**Comparison of Aneuploidy Detection to Somatic Mutation Detection**

For this study, we evaluated aneuploidy in plasma from 1348 healthy individuals and 883 cancer patients. We selected a cutoff (0.441) that produced 99% specificity in 1348 healthy individuals and calculated sensitivity on the 883 cancer patients. A detailed table of the aneuploidy results is included in Supplementary Data 4.

Cohen et al evaluates 812 plasma samples from healthy patients for somatic mutation detection. We selected an omega cutoff (1.77) that produced 99% specificity and calculated sensitivity on the same 883 cancer patients that were used in our study.

**Detection of cancer using a multi-analyte test**

Cohen et al demonstrates that combining somatic mutations and protein markers using a logistic regression model can predict cancer status of plasma samples. We compared whether aneuploidy could be integrated as an additional biomarker into the published framework as well as compared the predictive ability of a logistic regression model with aneuploidy and protein markers against the original logistic regression model that uses somatic mutations and protein markers.

In Cohen et al 812 plasma samples are from healthy people and 1005 are from cancer patients. Our study analyzed 1348 plasma samples from healthy people and 883 cancer patients. Of the 1348 healthy samples, only 248 overlapped with the original study. All 883 cancer samples were included in the original study. The sample demographic information was provided in Supplementary Data 5.

Using the original 812 healthy samples and the 883 cancer samples, we trained a logistic regression model and assessed performance using ten rounds of tenfold cross validation. A full list of samples and their biomarker values was provided in Supplementary Data 6. Because 564 of the original healthy samples were not analyzed for aneuploidy, we randomly sampled the list of scores from the 1348 normal samples and assigned each missing sample an aneuploidy value. Ten rounds of analysis were performed and each new round, we randomly sampled the collection of 1348 normal scores again to assign the 564 samples a new score.

To account for variations in the lower limits of detection across different experiments, we found the 90^th^ percentile feature value in the healthy training samples. We then found any feature value below that threshold and set all values to the 90^th^ percentile threshold. This transformation was done for all training and testing samples. This procedure was done for aneuploidy scores, somatic mutation scores, and protein concentrations. The 90^th^ percentile thresholds were listed in Supplementary Table 6. The results from each round and each cross validation fold for were included in Supplementary Data 7. The final feature coefficients from the logistic regression model were listed in Supplementary Table 6.

1.	Douville, C. et al. Detection of aneuploidy in patients with cancer through amplification of long interspersed nucleotide elements (LINEs). *Proceedings of the National Academy of Sciences* **115**, 1871-1876.

2.	Langmead, B. & Salzberg, S.L. Fast gapped-read alignment with Bowtie 2. *Nature methods* **9**, 357 (2012).

3.	Kinde, I., Wu, J., Papadopoulos, N., Kinzler, K.W. & Vogelstein, B. Detection and quantification of rare mutations with massively parallel sequencing. *Proceedings of the National Academy of Sciences* **108**, 9530-9535 (2011).

4.	Straver, R. et al. WISECONDOR: detection of fetal aberrations from shallow sequencing maternal plasma based on a within-sample comparison scheme. *Nucleic acids research* **42**, e31-e31.
